# Evaluation Of Effect Of Bacteria On The Surface Of The Dental Implant

**DOI:** 10.1101/2023.01.05.522965

**Authors:** S. Ajrish George, N.P. Muralidharan, Sahana S, Thiyaneswaran N, Subhashree R

**Affiliations:** Department of Implantology, Saveetha Dental College and hospitals; Department of microbiology, Saveetha Dental College and hospitals

**Keywords:** surface characterization, dental implant, bacterial action, SEM

## Abstract

**Background:** Dental Implantology is the most preferred treatment among patients who are opting for the replacement of their missing teeth. It is said that the success of the dental depends on the effectiveness of the dental implant bonding to the maxilla/mandible. This effectiveness depends on many factors. The microbial flora surrounding the dental implant also affects the integrity of the dental implant.

**Aim:** The study aims to analyze any effect bacteria on the surface of the implant.

**Materials and methods:** In the present study, 10 sterile dental implants were taken. Plaque samples were collected from patients with poor oral hygiene, who underwent dental implants and were under review for more than one month to six months. It was incubated with broth at 37 degrees Celcius. The surface changes in the dental implant were then observed in a scanning electron microscope.

**Results:** The SEM results show a significant difference in the surface texture of the implant. There were pits, small craters, and cracks on the surface. The sharpness of the screw thread had also been noticeably blunt.

**Conclusion:** Thus the study concludes that bacteria have a significant effect on the surface of the dental implant.

## INTRODUCTION

A dental implant is defined as A prosthetic device made of alloplastic material(s) implanted into the oral tissues beneath the mucosal or/and periosteal layer, and on/or within the bone to provide retention and support for a fixed or removable dental prosthesis; a substance that is placed into or/and upon the jaw bone to support a fixed or removable dental prosthesis.

The dental implant is a recent trend in dentistry. Implant survival is the main criterion for success and the majority of clinical studies show high success rates for dental implants. This is because of the biocompatibility of the titanium and its alloys. The biocompatibility of the commercially available pure titanium and its alloys is closely related to the surface properties. The composition of the protecting oxide film that is formed over the implant surface and the surface topography plays an important role in the success of the implant. The presence of 4 to 6 nm thick titanium oxide film that forms spontaneously when the surface of the implant is exposed to air or water which in turn is responsible for the excellent corrosion resistance of the implant. The surface chemistry of the implant might also influence other properties including adsorption of specific cell-binding proteins which is believed to depend both on surface energy and the sign and density of the surface charges in the implant. On the other hand, both the composition of the alloy and the manufacturing processes of the alloy will tend to influence the composition and the structure of the oxide film that is formed in the implant-bone interface. So this plays an important role in the biocompatibility properties of the dental implant. Surface contamination is believed to be important. This is because a fresh titanium oxide surface is very reactive toward both inorganic and organic contaminants

Biofilm formation on implant surfaces is similar in composition and mechanisms to that of the natural teeth but maybe additionally influenced by the special micro and macroscopic design features in the implant surfaces. The surface roughness of the implant has been reported to determine the shear strength of the implant-bone interface which is important for long-term fixation. But on the cell level and surface topography is known to influence the cell adhesion, morphology, proliferation, and differentiation of the cells and therefore has a major influence on the properties of the implant tissue interface. There are several methods for surface treatments that are used today on an industrial scale to achieve desired surface properties which include cleanliness, passivation, and specific topography

Exposed titanium is very easily colonized by bacteria which results in biofilm growth, corrosion pits, and subsequent deterioration of the mechanical performance of the dental implant.

The oral cavity contains thousands of species of microorganisms. Many of these microorganisms have oxidative and reductive properties on the metal surface. The prognosis of the treatment depends on the stability and the surface property of the metal.

The study aims to associate any relation of bacteria on the surface of the dental implant.

## MATERIALS AND METHODS

The study was done in department of implantology3 Saveetha Dental College, Chennai, India. The samples collected were from patients who underwent implant procedure from june 2021 to september 2021.

The plaque samples were collected from 10 included patients with poor oral hygiene and was preserved. Plaque samples were used on sterile implants to simulate the oral environment. The implants were placed individually in 10 different containers containing the plaque samples. The samples were incubated for 30 days at 37 degrees Celcius. To identify the specific organism, in this study a separate subculture was done. 10 microliters of the plaque sample were diluted with 2ml of saline and were agitated well. Then from the diluted plaque samples 10 microliters each was placed in nutrient agar, Macconkey agar, and blood agar, and streaking was done. After the inoculation of the plaque samples in the culture media, it was then incubated at 37°C overnight for microbial growth. There were observed microbial colonies in the culture media the next day. From the microbial colony present, a particular colony in each type was taken was smeared into a glass slide. Gram staining of the slide was done and was observed in the microscope. The species which was identified was then confirmed and noted separately. The dental implants which were incubated for 30 days were taken. The broth was drained from the Eppendorf. The implants were then washed in a good amount of distilled water. The implants were then placed in a new Eppendorf which was then labeled. The dental implants were then sterilized in an autoclave at 15lbs pressure(121°C) for 30 minutes. The sterilized dental implants were then dried in a hot air oven for 10-15 minutes. Then the dental implant was subjected to SEM analysis to analyze the surface changes. The exposed dental implants were compared to the unexposed sterile implants.

### Results

With three different media(nutrient agar media, McConkey agar media, and blood agar media) the organism in the plaque samples was cultured. From it, gram staining was done for each different colony obtained in the cultured plaque sample. 8 different organisms were identified in the gram staining. The plaque samples from which each organism was found in the gram staining were separately noted. The stock culture was done for each specific organism found. The bacterial colonies with different morphology isolated were lactobacillus species, beta-hemolytic streptococci, coagulase-negative streptococcus mutans, enterococcus, and bacillus species

The SEM observation was done in 1micometre magnifications. Thread diameter was measured. An increase of 1±0.03 μm in the thread diameter was observed in the exposed sterile implants when compared with the control unexposed implant threads. The sharpness of the implant screw threads was significantly reduced which was observed at about 0.5μm magnification

### Discussion

Bacterial infestation is one of the main causes of many failures in treatment procedures. Thought most of the time it cannot be avoided that there will be a bacterial-free environment because we live in harmony with the micro-organism. The oral cavity, it’s said to have more micro-organisms. Many of the microorganisms serve a purpose. But few of the microorganisms cause ill effects. Most of the time these microorganism is responsible for the failure of the treatment which causes secondary infection by invading the hard and soft tissue of the oral cavity. In implant dentistry, microorganisms play a vital role since if there is an infestation of the microorganisms then the osseointegration of the implant to the bone will be hampered leading to loss of bone, decrease in primary stability, and failure of the implant. So in the present study, the effects of this bacterial organism on the surface of dental implants are evaluated. In the present study, a scanning electron microscope is used as an imaging tool. The plaque samples were collected from 10 different persons to mimic the different oral condiction of different persons. The implants were incubated to allow the microrganim in the plaque to have a biofilm formation and if it affects the surface then we will be able to observe any surface changes(figure 1). The individual microorganism was identified using three media. The found organisms include lactobacillus species, beta-hemolytic streptococci, coagulase-negative streptococcus mutans, enterococcus, and bacillus species (figure 2). Pathogenicity of the each organism is different modes of action. In scanning electron microscopy, various changes were observed in the implant surface. An increase of 1±0.03 μm in the thread diameter was observed in the exposed sterile implants when compared with the control unexposed implant threads. The sharpness of the implant screw threads was significantly reduced which was observed. There were significant pits observed on the implant surface(figure 3). Further deep pits that were interconnected with fissures were observed. There was discontinuity of metal pattern with peaks of metal followed by a depression between the peaks is well observed. The microbial adherence to the metal surface is well observed. The individual microbial colony is present in various areas of the implant surface. Few groups of microbial colonies were also observed on different areas of the body of the implant. Several crater-like patterns were observed in various parts of the implant surface. These SEM findings were compared to a control implant(figure 4) which shows a regular pattern which is the stock pattern of the sterile implant. This shows a positive effect of surface damage caused by the microorganism. Further as a continuation of the study with more dental implant and more specificity to be added to the future undergoing study.

**Figure 1.**
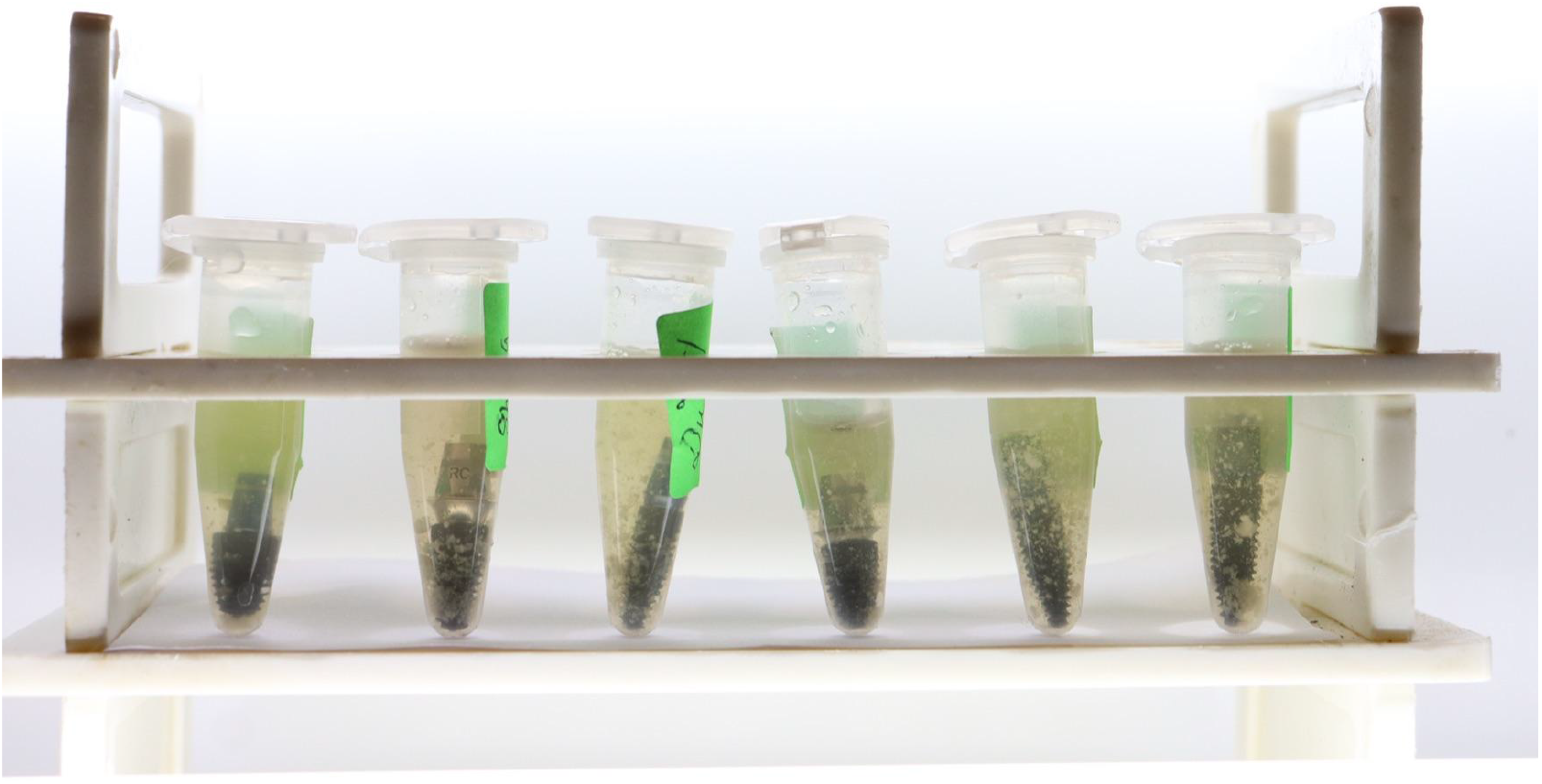
showing the incubation of the implant

**Figure 2.**
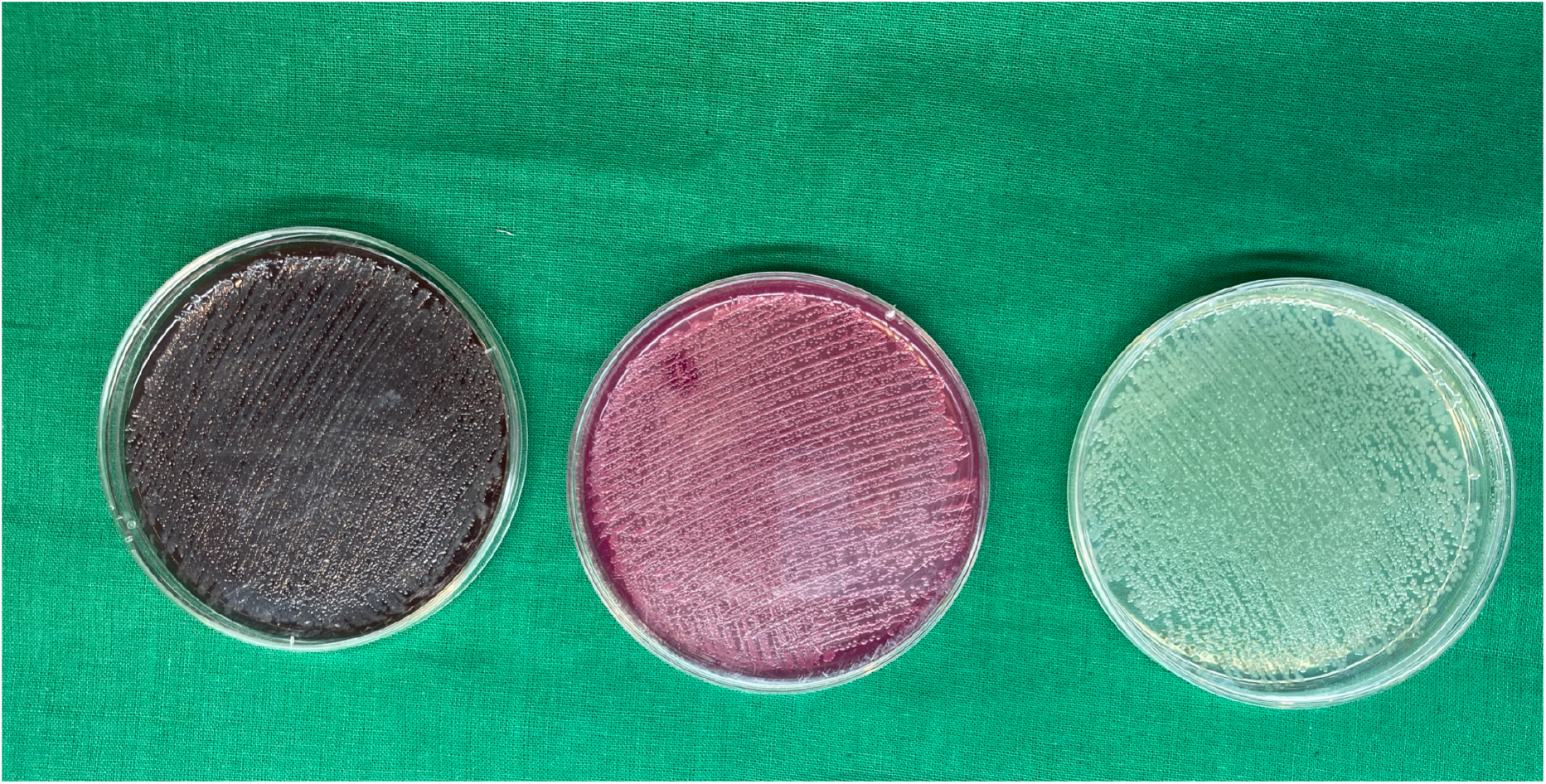
showing the individual bacterial organism

**Figure 3.**
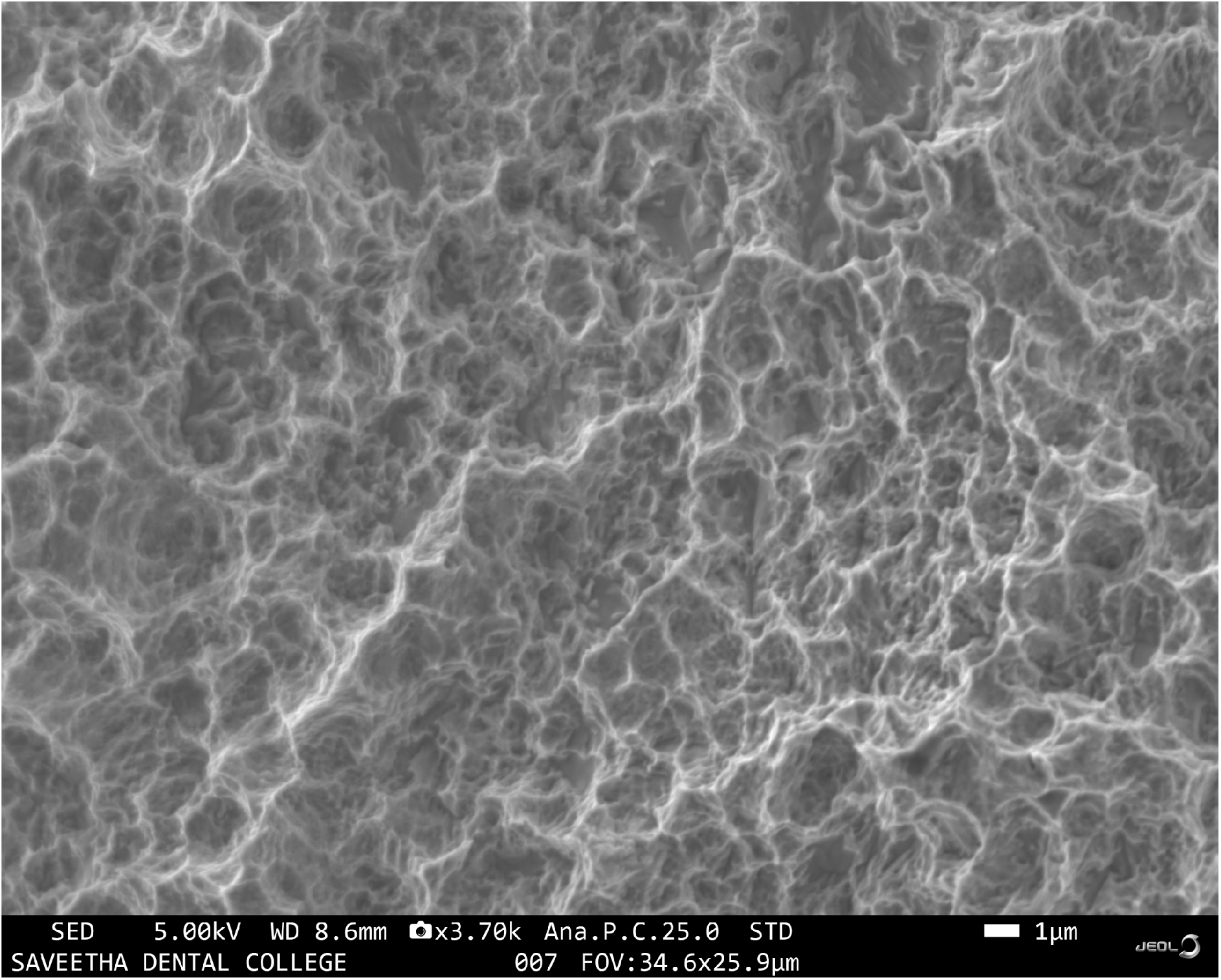
shows pre-incubation scanning electron microscopic image

**Figure 4.**
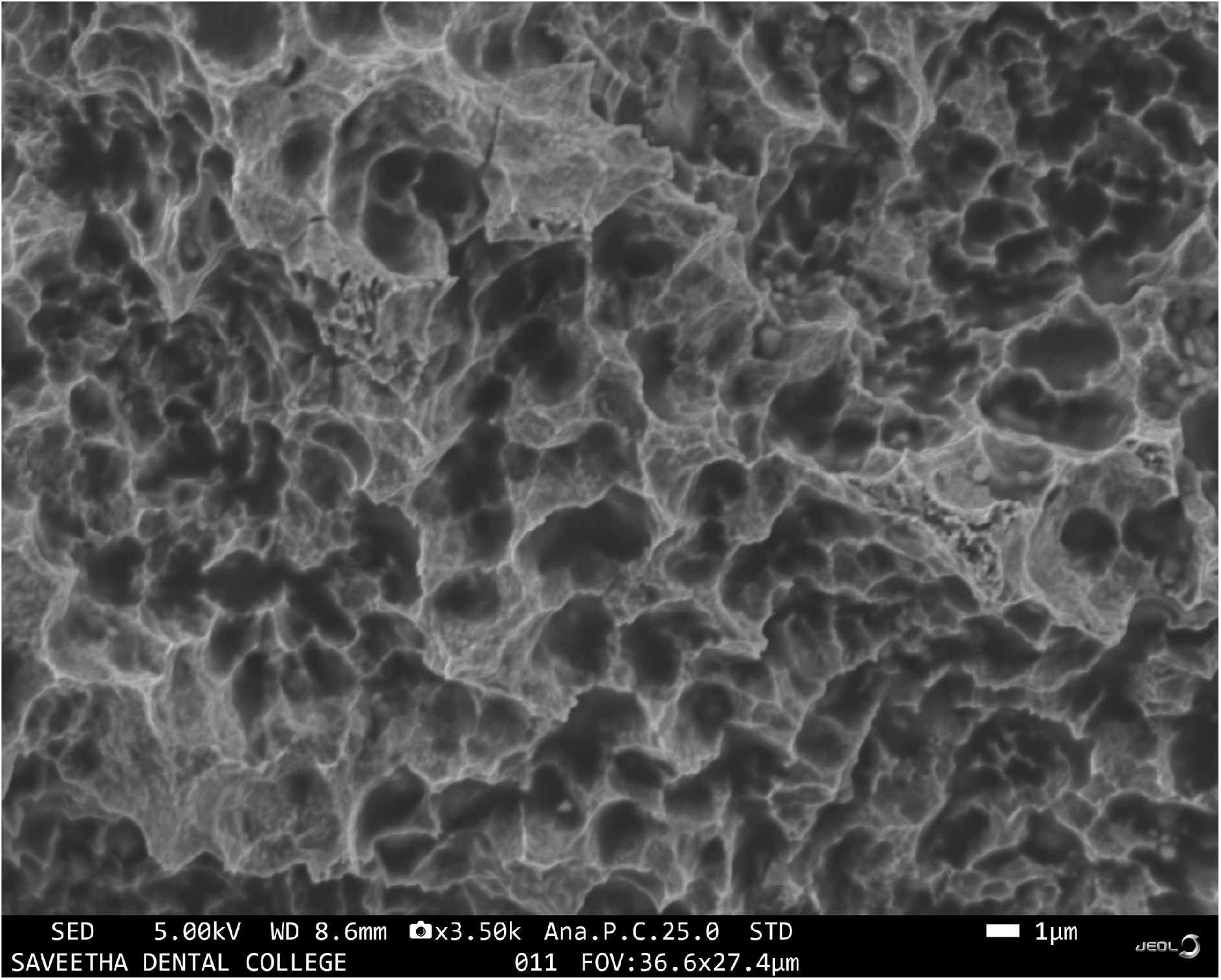
showing post-incubation scanning electron microscopic image

### Conclusion

Thus from the current study, the bacteria in the oral flora have an effect on the implant surface. This reveals the importance of maintenance of the teeth to prevent plaque and calculus accumulation as it can hamper the survival of the implant.

